# Undersampling techniques for non-linear chemical space visualization

**DOI:** 10.1101/2025.07.03.663077

**Authors:** Akash Surendran, Krisztina Zsigmond, Ramón Alain Miranda-Quintana

**Affiliations:** Quantum Theory Project and Department of Chemistry, University of Florida, Gainesville, Florida 32611, United States

## Abstract

The visualization of high-dimensional chemical space is a critical tool for understanding molecular diversity, structure–property relationships, and for guiding compound selection. However, the performance of non-linear dimensionality reduction (DR) techniques like t-Stochastic Neighborhood Embedding (t-SNE), Uniform Manifold Approximation and Projection (UMAP), and Generative Topographic Mapping (GTM) are often susceptible to the choice of hyperparameters, along with the high cost of their training for large datasets. In this study, we investigated the effect of undersampling methods on the choice of hyperparameter selection for these non-linear dimensionality reduction methods. Our results demonstrate that selecting small representative subsets of chemical data not only reduces computational costs associated with hyperparameter training but also serves as an innovative means to train non-linear DR methods, leading to projections that better preserve the local structure within the chemical space.

## Introduction

The chemical space represents the multidimensional landscape of already existing as well as theoretically possible molecules, enclosing the “intermolecular relationships” based on the structural and functional properties of these molecules.^1–6^ With advances in modern synthetic and computational fronts, this landscape is continuously expanding at a rapid rate, with estimates suggesting about 10^60^ molecules residing in the drug-like chemical space.^5^ These molecules are usually represented using vectors that span several hundreds or even thousands of dimensions, the simplest being binary fingerprints, where the presence and absence of a structural feature are represented by 1 and 0, respectively.^7–9^ Similarity indices are usually employed to quantify the similarity between a pair of molecules, with the Tanimoto similarity^10^ being the most widely used index for binary fingerprints. Mathematically, this function maps a pair of molecular fingerprints to a number between 0 and 1, with 0 and 1 representing completely dissimilar and identical molecules, respectively. The principle behind similarity-based searches is that “similar molecules display similar properties”.^11^

Typically, the average similarity of a library is calculated using the set of all pairwise similarities, which scales as O(*N* ^2^) with the number of molecules. To address this issue, our group developed iSIM,^12–16^ an extended similarity-based^17–21^ tool to calculate the average similarity of molecular libraries in O(*N*). The principle behind iSIM is the simultaneous comparison of multiple molecules, and it has been shown that the results are consistent with pairwise similarity across both binary fingerprints and real-valued descriptors. Coupled with complementary similarity, this method also supports advanced sampling of molecular libraries to target specific sectors of the vast chemical space, improving the coverage and representation of the selected compound sets.

Following the identification of promising regions, a crucial next step is the visualization of the chemical space to interpret the complex structure-property relationships between molecules. Because the usual high-dimensional representation of molecules is not interpretable to the human eye, dimensionality reduction (DR) methods are usually employed to project these fingerprints to two or three dimensions.^22–24^ Principal Component Analysis (PCA)^25,26^ remains a workhorse technique for chemical space visualization, which works by identifying orthogonal axes of maximum variance in the descriptor space and projecting them along these axes, preserving the global data structure. However, this comes at the cost of poor local neighborhood preservation, especially when the chemical space is highly non-linear. In a recent study by Orlov et al.,^23^ it was shown that non-linear methods such as t-Stochastic Neighborhood Embedding (t-SNE), Uniform Manifold Approximation and Projection (UMAP), and Generative Topographic Mapping (GTM) provide far superior performance in preserving the local structure of the underlying chemical space, which mostly consists of small organic molecular compounds. Similarly, Takacs et al.^27^ investigated the application of self-organizing maps (SOMs) to visualize and analyze regions of drug-like chemical space that remain heavily underrepresented. This method maps the high-dimensional chemical space onto a two-dimensional grid, preserving the topological features and highlighting the property clusters. By transforming multidimensional chemical data into intuitive graphical representations, these methods can enhance the virtual screening of large molecular libraries, structure-activity relationships (SAR), library design, and synthesis of drug-like molecules.^4,5,28^

Non-linear manifold-based dimensionality reduction methods on one hand provide key insights into the local distribution of data in molecular libraries but require extensive computational resources. The hyperparameter set plays a key role in the efficiency of these methods, and training these models on larger datasets can be challenging. In this study, we attempt to address this problem by training three non-linear models, t-SNE, UMAP, and GTM on 30 CHEMBL datasets^29^ by implementing five different sampling methods using iSIM, but instead training the models on these smaller datasets. A comparison will be performed to determine whether the choice of sampling can help improve the quality of projections performed by these three DR methods. The performance of the projections will be evaluated on several local and global neighborhood preservation metrics^23,30^ (discussed in the next section), and the results will be compared across different DR methods as well as within each DR method across different training samples used. The key question to be answered in this manuscript is thus: what is the impact of robust, deterministic, undersampling methods on the selection of training sets for non-linear DR techniques in chemical space visualization.

## Methodology and Workflow

### Generation of Fingerprints

A brief visual representation of the workflow is provided in Figure 1. CHEMBL datasets for 30 different macromolecular targets containing between 615 and 3657 molecules in SMILES strings were selected from the study by van Tilborg et al.^29^ For the larger benchmarking study, a subset of 50000 molecules was created from the mcule natural products dataset.^31^ Binary Morgan fingerprints were generated using the RDKit module in Python.^8,32^ In this representation, molecules are represented as fixed-length binary vectors, and the popular Extended-Connectivity Fingerprints (ECFP) with a radius of 2 and 1024 bit length, also known as the ECFP4 representation, was employed. Although the choice of molecular representation mainly depends on the focus of the study, here we are particularly interested in the local chemical space mapped by ECFP4 fingerprints, since mapping faithful latent representations is important for structure-activity relationship studies. Hence, we will focus on the local neighborhood preservation between the fingerprint space and latent space rather than the entire global structure of the dataset.

**Figure 1:**
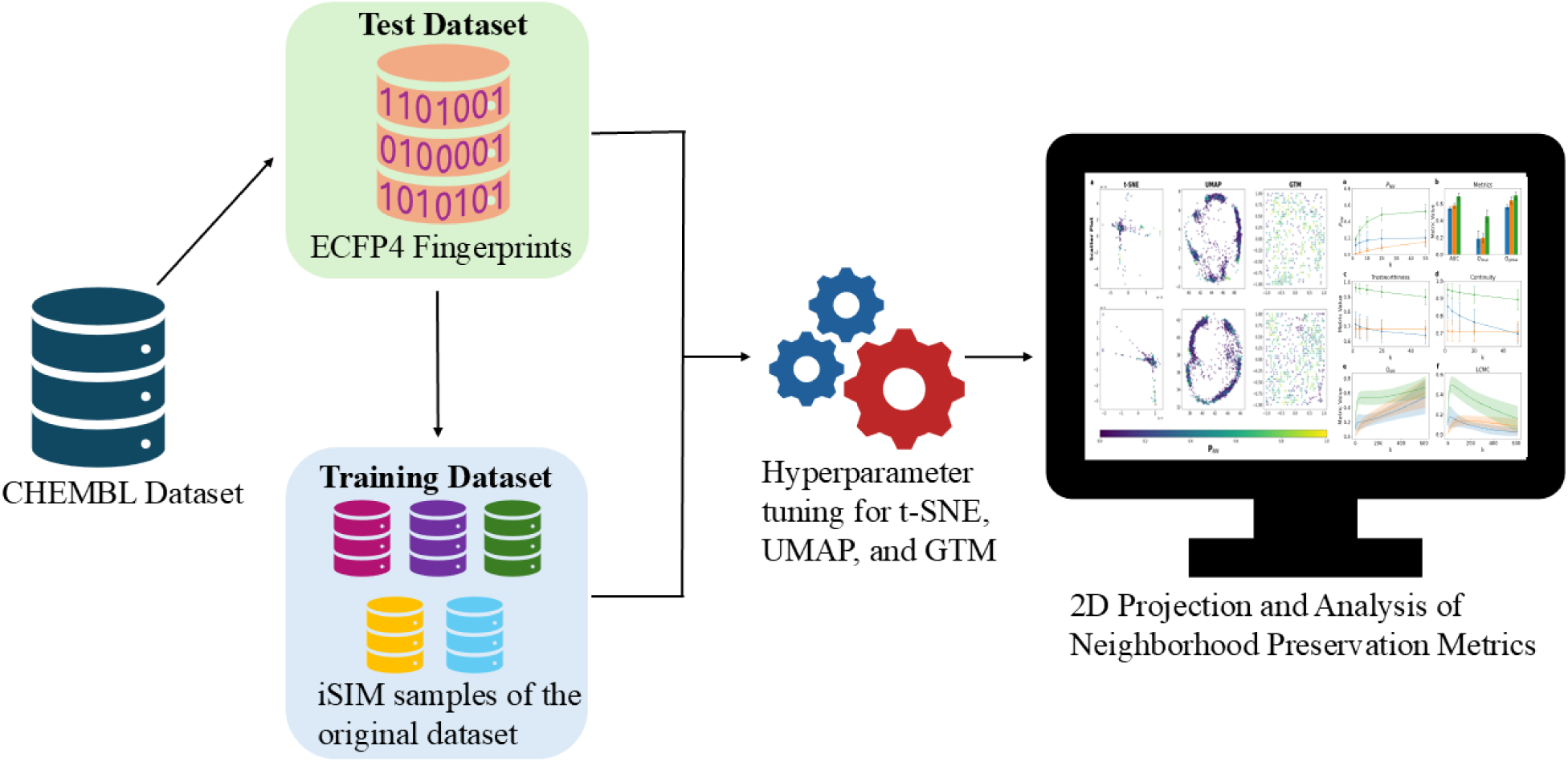
Brief workflow: 30 CHEMBL datasets curated for 30 different macromolecular targets were selected for this study. Morgan fingerprints were generated for each dataset, and samples (10% and 20% of the original dataset size) were generated using iSIM complementary similarity sampling methods. Non-linear DR methods were trained using each sample, and the corresponding original dataset was projected in 2D. Neighborhood Preservation Metrics were analyzed to compare the quality of these projections for chemical space visualization.

### iSIM complementary similarity based sampling of libraries

iSIM is a novel computational method that efficiently calculates the average similarity between molecular sets in linear time scaling (O(*N*)), bypassing the quadratic (O(*N* ^2^)) scaling of traditional pairwise approaches. It works by aggregating bit- or descriptor-level statistics across the entire set to calculate the average similarity of the chemical library. For example, to calculate the instant Tanimoto similarity (*iT*) for a set of *N* molecules represented by *M* −dimensional binary fingerprints, the first step is to arrange all these vectors in a matrix with each row representing a molecule, that is, an *N* × *M* matrix. Then, a 1 × *N* matrix corresponding to the sum of the individual columns of the previous matrix is constructed, and each element of this matrix is represented by *k_q_*, *q* = 1, 2*,…, M*. Using this, *iT* can be calculated as

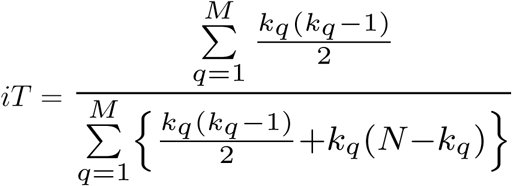

After fingerprint generation, the datasets were sampled using complementary similarity-based sampling tools in the iSIM module.^12^ The key idea behind complementary similarity is “how well a molecule represents others.” One molecule was removed, and the iSIM for the remaining set was calculated. A higher complementary similarity corresponds to low-density regions or lower similarity to the rest of the set, whereas the opposite is true for lower complementary similarity. Similarities were calculated using the Tanimoto similarity index, which is a widely used similarity index for binary data, and the molecules were ranked according to increasing complementary similarity. This can then be used to target specific regions of chemical space by providing a means to sample the set in *O*(*N*) complexity. Using the tools in our group’s GitHub https://github.com/mqcomplab/iSIM/tree/main, we sampled 10 and 20% of each dataset by applying the following five sampling methods:

- Medoid sampling: M (10 or 20% of the total molecules in the set) molecules with lowest complementary similarity
- Outlier sampling: M molecules with highest complementary similarity
- Extremes sampling: M/2 molecules with highest complementary similarity and M/2 molecules with the lowest complementary similarity.
- Stratified sampling: After the molecules are arranged from lowest to highest complementary similarity, this set is divided into b strata of equal size (containing same number of molecules) and molecules with the lowest complementary similarity are selected from each stratum until M molecules are chosen.
- Quota sampling: The range of complementary similarity (max-min) is divided into b strata of equal range which is 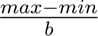. Note that this time the range of complementary similarity in each stratum is same but not necessarily the number of molecules. Molecules are then chosen till M is reached.

The key difference between quota and stratified sampling lies in the way the set is divided. For stratified sampling, the strata are created such that they contain the same number of molecules but not necessarily the same comp sim range, whereas for quota sampling, the strata are created such that they correspond to equal comp sim range but not necessarily the same number of molecules. A visual representation of all five sampling methods is given in Figure 2 for CHEMBL234, the largest dataset in this study, which contains 3657 molecules. Medoid, outlier and extremes target specific regions of the chemical space, while stratified and quota samples target diverse regions of the underlying chemical space.

**Figure 2:**
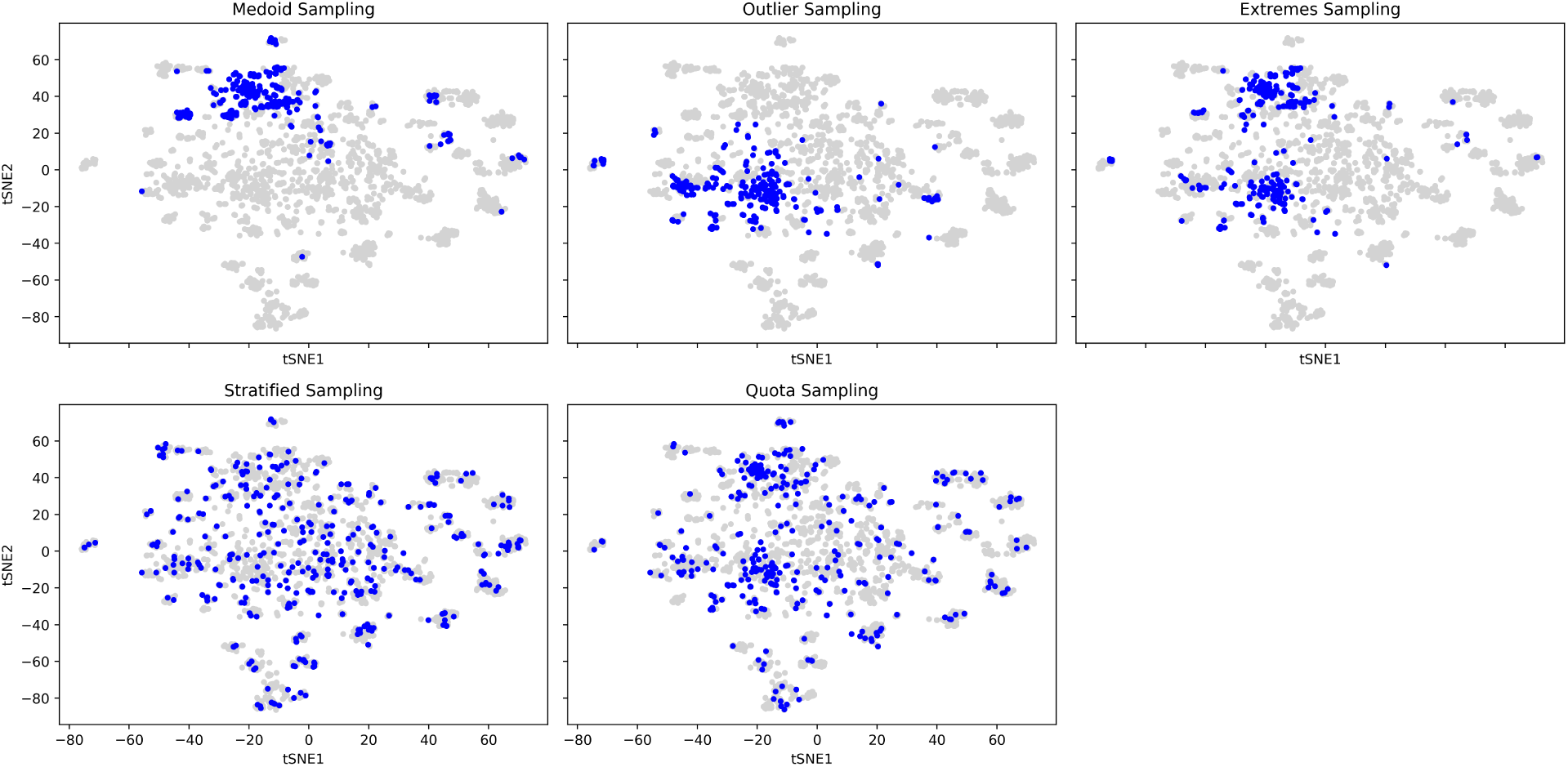
t-SNE plots for CHEMBL234 by sampling 10% of the data. Blue portions depict the sampled molecules and grey portions represent the projection of the entire dataset. The former three methods target specific portions but the latter two target diverse regions of the chemical space.

### Hyperparameter tuning of non-linear DR methods

The next step involved hyperparameter tuning for the three non-linear dimensionality reduction methods: t-SNE, UMAP, and GTM. DR models and metrics were chosen from the GitHub repository of the authors of.^23^ The t-SNE algorithm was implemented using the OpenTSNE Python library, the UMAP algorithm using the umap-learn library, and the GTM using the Incremental GTM of Gaspar et al.^33^ The set of hyperparameters was exactly the same as that in, ^23^ and a grid search was adopted to tune the best set of hyperparameters. For t-SNE, perplexity values were selected from [1, 2, 4, 8, 16, 32, 64, 128] and exaggeration from [1, 2, 3, 4, 5, 6, 8, 16, 32], learning rate as the default of OpenTSNE and Fast Fourier Transform accelerated interpolation was used for gradient calculation. For UMAP, the nearest neighbors (n neighbors) were chosen from [2, 4, 6, 8, 16, 32, 64, 128, 256], and the minimal distance (min dist) was chosen from [0.0, 0.1, 0.2, 0.3, 0.4, 0.6, 0.8, 0.99]. For GTM, the number of nodes was selected from [225, 625, 1600], the number of basis functions was set to [100, 400, 1225], the regularization coefficients were set to [1, 10, 100], and the basis widths were set to [0.1, 0.4, 0.8, 1.2]. The scoring was based on the percentage of preserved nearest 20 neighbors in the high-dimensional fingerprint space.

### Neighborhood Preservation Score

In a pivotal contribution, Orlov et al.^23^ recently highlighted the relevance of quantifying the degree to which different DR representations could retain information about local environments compared to the original high-dimensional fingerprint space. A combination of local and global preservation metrics was used to evaluate the efficiency of the neighborhood preservation. The first step involves finding the first *k* neighbors of a molecule in the “ambient” or 1024-dimensional fingerprint space based on the Tanimoto similarity. The same step is then repeated in the “latent” or low-dimensional projection space, but using Euclidean distances, since the Tanimoto distance is not a metric for real-valued vectors.^34^ The number of nearest preserved neighbors can be evaluated using the neighborhood preservation score *P_NN_* (*k*):

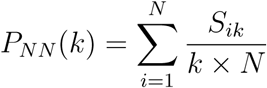

where k represents the number of neighbors considered and *S_ik_* represents the overlap or shared k-nearest neighbors of the *N* compounds in latent and ambient spaces. Additional local and global preservation metrics can be evaluated by constructing the co-ranking matrix Q, which is discussed in the next section.

### Co-ranking Matrix Q and Neighborhood preservation metrics

A matrix can be constructed in both the ambient and latent spaces by evaluating and ranking the pairwise distances. This matrix is called the co-ranking matrix Q. The elements of the co-ranking matrix *Q_kl_* count the number of cases where samples of rank k in the ambient space become rank l in the latent space and are diagonal in the ideal case. This matrix can also be used to calculate the following DR metrics.

1. **Co-k Nearest Neighbor size**: Represents the number of points within k-nearest neighbors before and after DR, giving the total number of mild intrusions and exclusions:

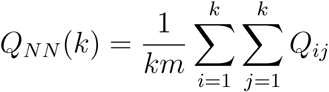

where m represents the total number of neighbors, and k is the number of neighbors considered for the hit calculation. The factor 1*/km* is a normalization factor that keeps the values in [0, 1] range. Here, mild intrusions refer to elements with *k < l* and correspond to far-away points pulled closer by the DR. Mild exclusions refer to points with *k > l* and correspond to points that are pushed away by DR. Ideally, *Q_NN_*is 1, which means no change in the order of neighbors after DR.

1. **Area under** *Q_NN_* **curve:** A single metric irrelevant of k to calculate the global neighborhood preservation based on the *Q_NN_* curve:

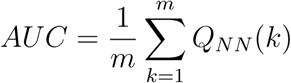

1. **Local continuity meta criterion (LCMC):** As an alternative to *Q_NN_*, LCMC removes the number of neighbors 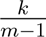 from *Q_NN_*:

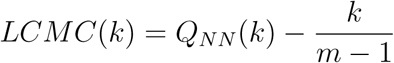

The metric favors local neighborhood preservation compared to *Q_NN_* by ensuring a larger penalty for a higher *k* value. *k* value corresponding to the maximum of *LCMC*(*k*), *k_max_* is

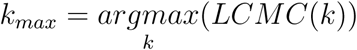

1. **Local and Global Property metrics:** Calculated from the *Q_NN_* curve, *Q_local_* and *Q_global_* correspond to the local and global neighborhood preservation metrics. Given the *k_max_* value previously defined, these metrics can be calculated as follows:

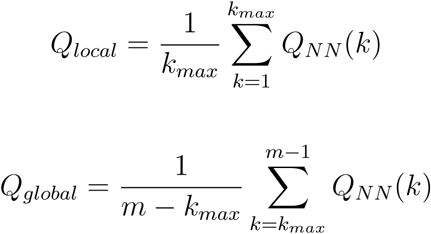

*Q_local_* is usually preferred over *Q_global_* because the immediate neighbors of a compound are much more significant in cheminformatics studies.

**5. Trustworthiness and Continuity:** Trustworthiness and continuity are used to account the errors due to hard-intrusions and hard-extrusions respectively.

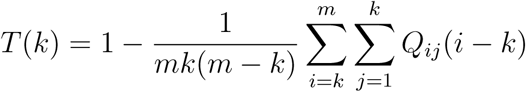

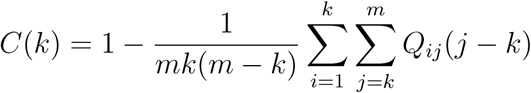

The DR methods and neighborhood metrics were generated with the help of software from Varnek’s group and the code is available at https://github.com/AxelRolov/cdr_bench.

## Results and discussions

### Benchmarking on 30 ChEMBL datasets

The grid search was performed as follows: each of the 30 ChEMBL datasets was first sampled to obtain 10 and 20% of the data using the iSIM sampling tools corresponding to the five sampling methods previously discussed. These sampled sets were then used to perform hyperparameter tuning for the three non-linear DR methods: t-SNE, UMAP, and GTM. Because these methods are susceptible to the choice of hyperparameters and the intrinsic structure of the data, there is no universal hyperparameter configuration, and the best selection of hyperparameters will always vary with the chosen datasets. For example, t-SNE was applied to published single-cell RNA-seq datasets to study the perplexity trade-offs between the local/global structure to show that although a high perplexity value (∼ 1% of sample size) can improve the global structure of the projected data, it comes at the cost of degraded local structures.^35^ Despite the acceleration of gradient calculations provided by FFT and interpolation, t-SNE is still typically slower than PCA and UMAP (for the order of ∼ 10^4^ datapoints), making the hyperparameter tuning process costly for large datasets. In our computation, the complete training and testing process of UMAP took the least time, 33s to train on 10% of the CHEMBL204 dataset (sample size = 274 molecules) and then project the entire dataset (containing 2754 molecules). On the other hand, GTM took the most time, about 12 minutes to do the same, followed closely by t-SNE which took about 11 minutes and 30s.

In a previous work by the group, ^12^ it was shown that iSIM achieves linear scaling (O(*N*)) in dataset sampling and when about 10% of the datasets were selected, quota and stratified sampling stand out in closely matching the average similarity of the entire dataset, achieving better coverage of the underlying chemical space. This is also clear from the visual representation of these sampling methods in Figure 2. Although the global data structure is better captured by quota and stratified samples, the main interest of this article is if the choice of sampling could influence the local structure after DR. A scatter plot of these three methods trained on the five sampling methods on one of the datasets used in the study (CHEMBL237) is shown in Figure 3. The structure of projections shows significantly larger variations on t-SNE trained across different sampling methods to than those for UMAP and GTM. This indicates that t-SNE is much more sensitive to the choice of hyperparameters than the other two DR methods. A larger portion of bright points in the GTM heatmap indicates superior local neighborhood preservation with the method trained on quota and stratified samples showing a larger number of bright points.

**Figure 3:**
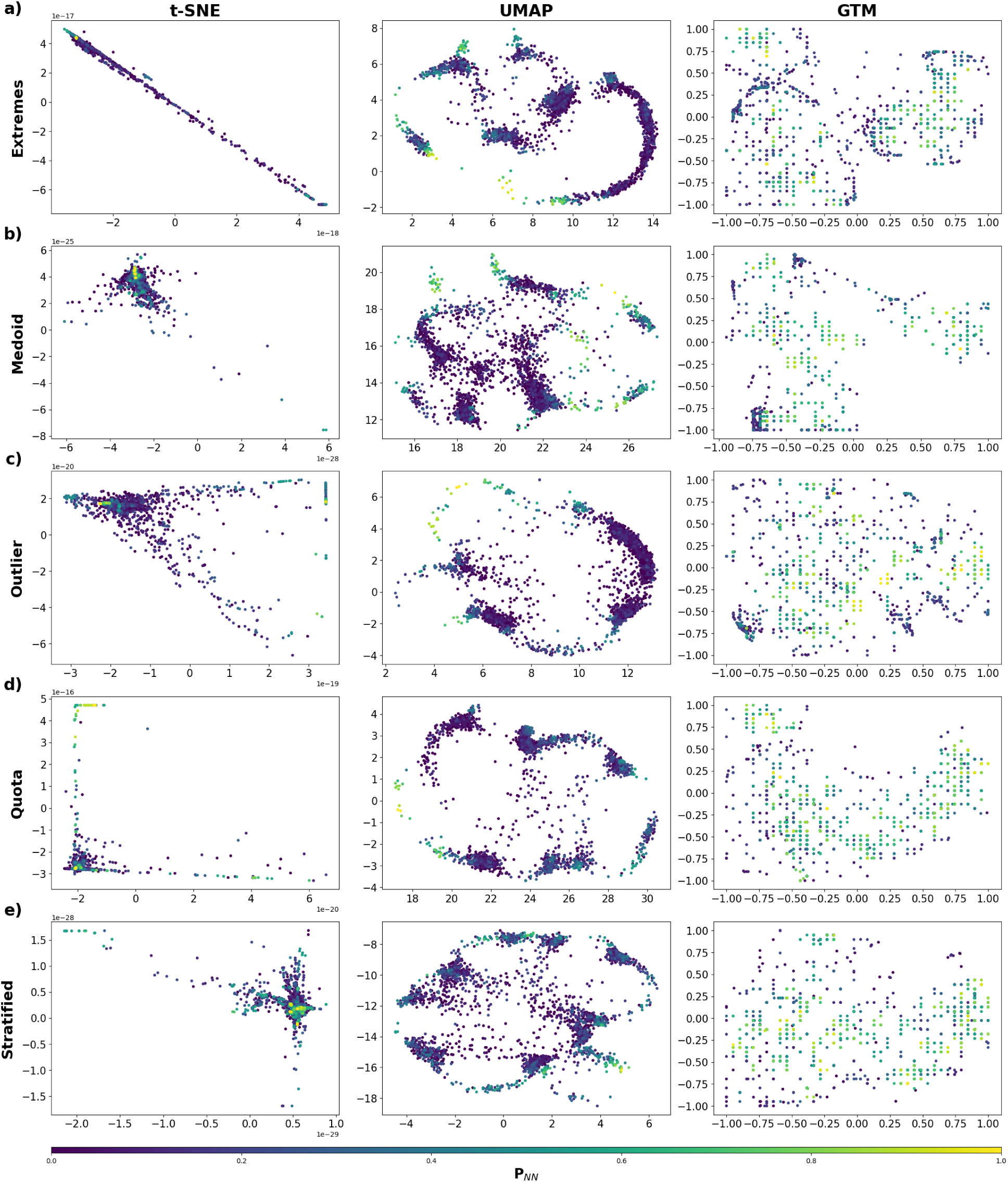
t-SNE, UMAP and GTM projections of CHEMBL237 dataset trained on 20% samples by Extremes, Medoid, Outlier, Quota and Stratified sampling. The abundance of lighter shades in the heatmap of GTM indicates a superior local neighborhood preservation. t-SNE on the other hand, is more sensitive to the choice of sampling.

To get a quantitative understanding of the quality of projections, the performance of these five sampling methods is evaluated based on the neighborhood metrics described in the previous section. The key idea is to sample datasets up to one or two orders of magnitude below the actual size, capturing as much information as possible about the distribution of the data, which further boosts the optimum hyperparameter selection at reduced computational costs. Considering the size of smaller datasets, the sample size was chosen to be at least 10%, to have a greater number of molecules than the minimum number of strata, which was set to 50 by default. The hyperparameter scoring is based on the percentage of the first 20 neighbors preserved in the latent space, according to the code developed for.^23^ Because we are interested in the local neighborhood of a molecule rather than the global structure of the entire dataset, special focus is placed on metrics such as *PNN* (*k*), *Q_local_*, Trustworthiness and Continuity, as the presence of similar structural features or functional groups governs properties.

A plot of these metrics against the number of neighbors *k* is given in Figure 4 and are tabulated in Table 1 (the values correspond to *k_hit_* = 20). *AUC* and *Q_global_*describe the global neighborhood preservation quality of the DR method, while *Q_local_*, PNN, trustworthiness, and continuity focus on local neighborhood preservation. Although the performance of t-SNE and UMAP was consistent across all five sampling methods, there were key differences in the performance of GTM. First, while comparing the local structure preservation metrics across these three methods, it can be seen that GTM offers a far superior performance to t-SNE and UMAP, while the global metrics, such as AUC and *Q_global_* do not show a significant difference. The percentage of preserved nearest 20 neighbors (PNN) was less than 20% for t-SNE and UMAP, while the number was as high as 48% on average for GTM across all 30 CHEMBL datasets. This value was as high as over half when the first 50 neighbors were considered. This hints at the far superior performance of GTM over the other two methods while offering a similar global structure.

**Figure 4:**
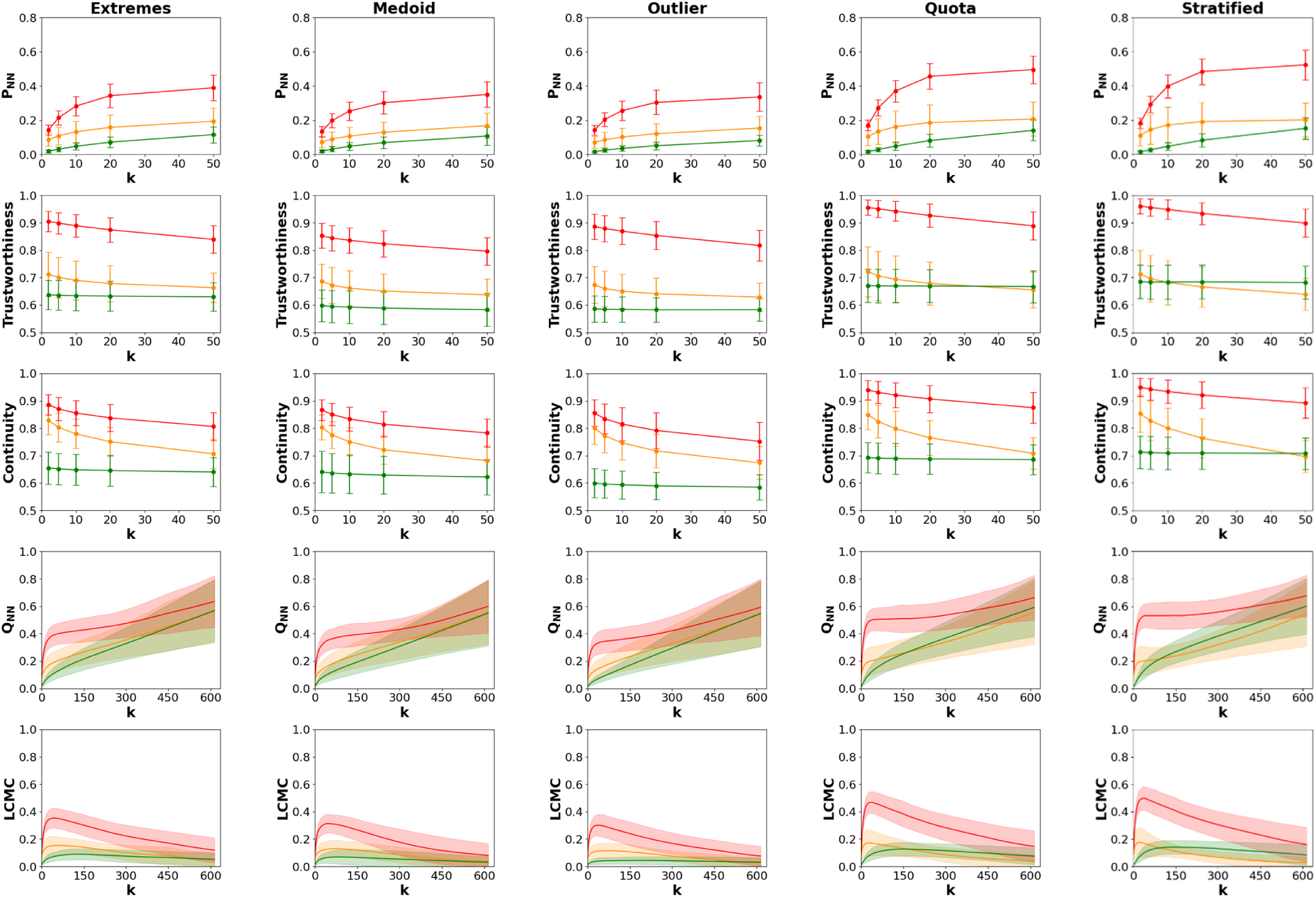
DR metrics plot in the order Extremes, Medoid, Outlier, Quota and Stratified. The color scheme is as follows: t-SNE: Yellow, UMAP: Green and GTM: Red. The shaded regions represent the standard deviation across datasets.

**Table 1:**
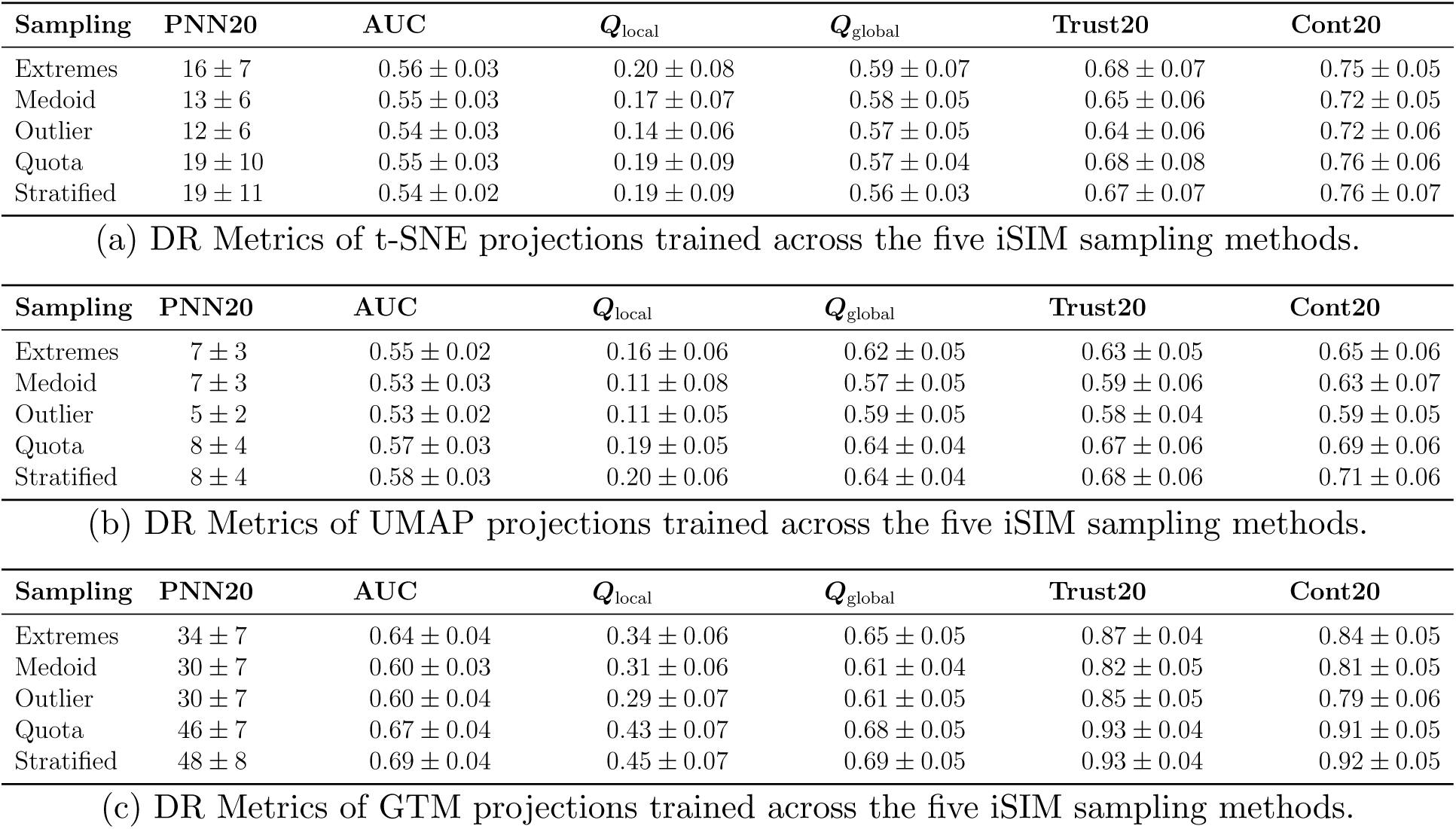
Embedding quality of neighborhood preservation metrics across the five sampling methods for (a) t-SNE, (b) UMAP and (c) GTM. While the global preservation metrics do not show a large variation across DR methods, the local preservation is consistently better in GTM. GTM trained on quota and stratified samples show enhanced local metric preservation while the performance of t-SNE and UMAP are similar across all sampling methods.

Given the rapidly growing amount of chemical data, a central problem in cheminformatics is to train Machine Learning models with minimum input to reproduce maximum information about a given dataset. Hyperparameter selection is key for the performance of non-linear DR methods and often computationally expensive given their non-linear scaling with data size, and hence, we address this problem by asking whether a smaller sampling of these datasets could provide a suitable set of hyperparameters and hence a better quality of projections. The neighborhood preservation metrics were compared within each DR method, and it was observed that although these metrics remained consistent across all sampling methods for t-SNE and UMAP, there were stark differences in the quality of projections in GTM. While *P_NN_* shows an increasing trend with the number of neighbors k across all DR methods, GTM trained on quota and stratified sampling of datasets displayed an improved performance, preserving as high as over half of the nearest 50 neighbors in the latent space. Trustworthiness metric displays significantly better values in GTM, but more importantly achieves an average value of 93% when GTM is trained on Quota and Stratified sampling. Even for *k* = 50, the metric shows a value above 90% while other training samples show about 80% and an even lower number (between 60 and 70%) for t-SNE or UMAP. A similar trend is also observed for the Continuity metric. Note that Trustworthiness and Continuity display the errors due to Hard-intrusions and hard-extrusions respectively. A high value for both metrics indicates that the DR method minimizes false positives (trustworthiness) and false negatives (continuity) in local relationships.

*LCMC*, which is a local preservation-focused alternative to *Q_NN_*, peaks at a higher value for quota and stratified training samples, leading to a better *Q_local_* score, an important local preservation metric. *Q_local_*, in general shows a relatively better performance even for t-SNE and UMAP, although, the differences were more profound when comparing within GTM, with average values as high as 0.45. A key reason for the superior performance of GTM compared to other DR methods is its probabilistic grid structure, which preserves the local topology better. Coupled with efficient iSIM sampling methods like quota and stratified sampling which provide a better description of the data distribution, hyperparameter selection is improved. Even with a small fraction of data (approximately 10-20%), the set of hyperparameters corresponding to the intrinsic distribution in the original data set can be found at a highly reduced cost, leading to a better and more accurate low-dimensional projection of the data. Sampling methods covering diverse chemical regions outperform extremes-focused approaches. Another key observation is that stratified/quota sample trained projections lead *Q_global_* by a small margin (approximately ≤ 0.08) across methods, indicating that global structure is less sensitive to sampling than the local structure. A similar trend can also be observed in the case of AUC.

### Benchmarking for a 50k mcule natural products subset

To evaluate whether the undersampling-guided hyperparameter selection generalizes to higher scales, we extended this benchmarking pipeline to a 50,000-molecule subset of mcule natural products which is roughly one to two orders of magnitude larger than the datasets used in the first study. Following the same procedure, each of the three non-linear DR methods (t-SNE, UMAP and GTM) were trained on a 10% sample (∼ 5000 molecules) drawn using each of the five iSIM complementary similarity based sampling strategies. Critically, PCA was introduced as a linear hyperparameter-independent dimensionality reduction baseline in this extended benchmark, providing an important reference point to disentangle method family from sampling effect. The full 50k dataset was then projected using the best hyperparameter settings identified from each training sample, and the resulting embeddings were evaluated across the same five neighborhood preservation metrics used previously: the nearest-neighbor preservation score *P_NN_*, Trustworthiness, Continuity, *Q_NN_*, and LCMC.

The results of the benchmarking are given in Figure 5. The neighborhood overlap score *P_NN_* reveals a striking reversal relative to the smaller dataset benchmarks: t-SNE (yellow) is now the top-performing method across all five sampling conditions and all values of neighbors considered k followed by GTM, UMAP and then PCA which exhibits a poor *P_NN_* score, barely exceeding 5%. All the methods show less than 20% overlap for the nearest neighbors considered for the former three sampling methods, whereas the *P_NN_* scores for t-SNE get as high as about 31% for this neighborhood size showing significant improvement for the case of stratified sampled configurations.

**Figure 5:**
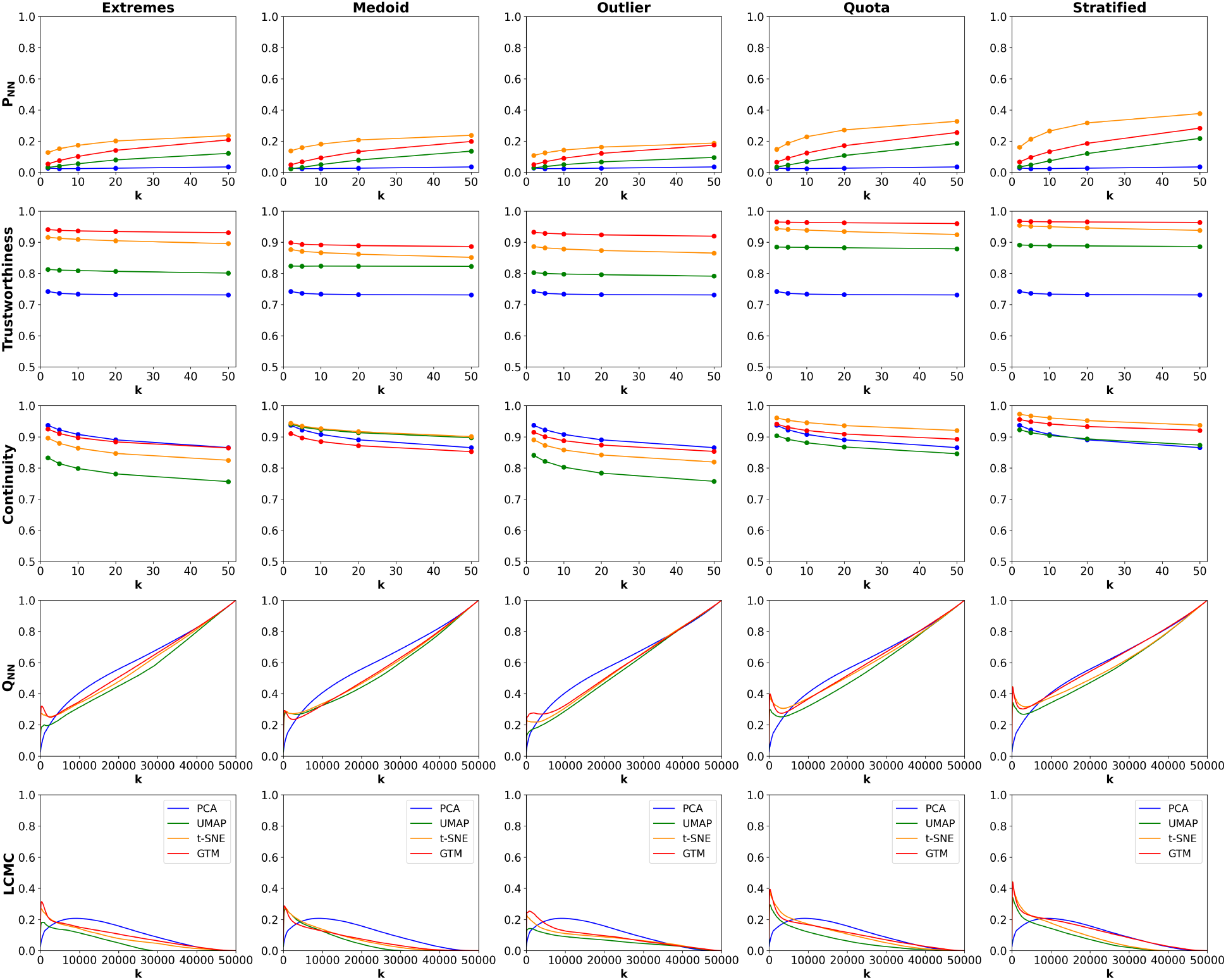
DR metrics plot for the 50k mcule natural product subset in the order Extremes, Medoid, Outlier, Quota and Stratified. The color scheme is as follows: PCA: Blue, t-SNE: Yellow, UMAP: Green and GTM: Red.

At the scale of 50000 molecules, the 10% training set is significantly larger than those used in the original benchmarking with much smaller datasets. The richer and more chemically diverse training sample appears to directly benefit t-SNE’s perplexity optimization: with more molecules in the training pool, the grid search identifies perplexity values that better capture the true local density of the full chemical space. GTM follows the trend in showing a similar improvement for the latter two sampling strategies, though it does not show the same proportional gain, possibly because its fixed grid resolution becomes a limiting factor as the dataset size grows. The number of latent grid nodes may not scale as naturally with dataset density of t-SNE’s perplexity hyperparameter.

UMAP relies on a local-global tradeoff and is known to prioritize the preservation of the broad topological skeleton of a dataset over fine-grained local neighborhoods, and this tendency appears to become even more pronounced as dataset size increases. At 50k molecules, the chemical space contains significantly more overlapping local dense regions than in smaller benchmarks, and even when optimized by grid search on 10% sample, UMAP may not adequately capture the heterogeneity of local density across the full dataset. The result is a systematic underestimation of the true local neighborhoods in the 2D projection, reflected in the persistently low *P_NN_* values. PCA, predictably, performs the worst because the method assumes linearity and relies on global variance maximization which is essentially blind to the local structure, and its projections compress the intricate local topology of chemical space into a low-dimensional linear subspace that discards most of the neighborhood information.

Trustworthiness also shows a similar trend where all three non-linear DR methods show elevated values for quota and stratified sampling, whereas PCA, due to its poor neighborhood information, exhibits significantly lower values. GTM maintains a slight edge in trustworthiness, while t-SNE’s higher *P_NN_* is accompanied by marginally lower trustworthiness values. This distinction is meaningful: t-SNE’s aggressive local clustering recovers a larger fraction of true neighbors at the expense of some false positives whereas GTM’s probabilistic grid on the other hand imposes a softer topology that suppresses hard-intrusions even at scale, leading to a slightly more reliable map. Unlike non-linear DR methods, PCA simply maps the data into directions of maximum variance irrespective of local topology, producing mediocre trustworthiness values.

The global neighborhood preservation scores like AUC and *Q_global_* show similar trends as in the previous results, see Table 2, where the choice of sampling appears to make little to no difference for these metrics and the trend is consistent across all DR methods. These results carry several important implications for undersampling-guided DR at large chemical dataset scale. Most importantly, the performance hierarchy is not fixed across dataset size as t-SNE, which showed significantly lower local preservation than GTM on the smaller datasets, emerged as the leading method at 50k molecules. This suggests that the relative advantage of each DR method is dataset-size dependent, and that the richer information available in a 10% sample of a large dataset particularly benefits t-SNE’s hyperparameter optimization. GTM remains the more trustworthy method in a strict false-positive sense, and the choice between the two should be guided by whether the application demands maximum neighbor recall (favoring t-SNE) or maximum projection reliability (favoring GTM). PCA’s near-zero performance on all local metrics at this scale delivers a clear message: for chemical libraries of this size and diversity, linear dimensionality reduction is not a viable tool for local neighborhood visualization and should be restricted to use cases where global variance structure is the primary interest, such as identifying broad chemical series or detecting outlier libraries.

**Table 2:**
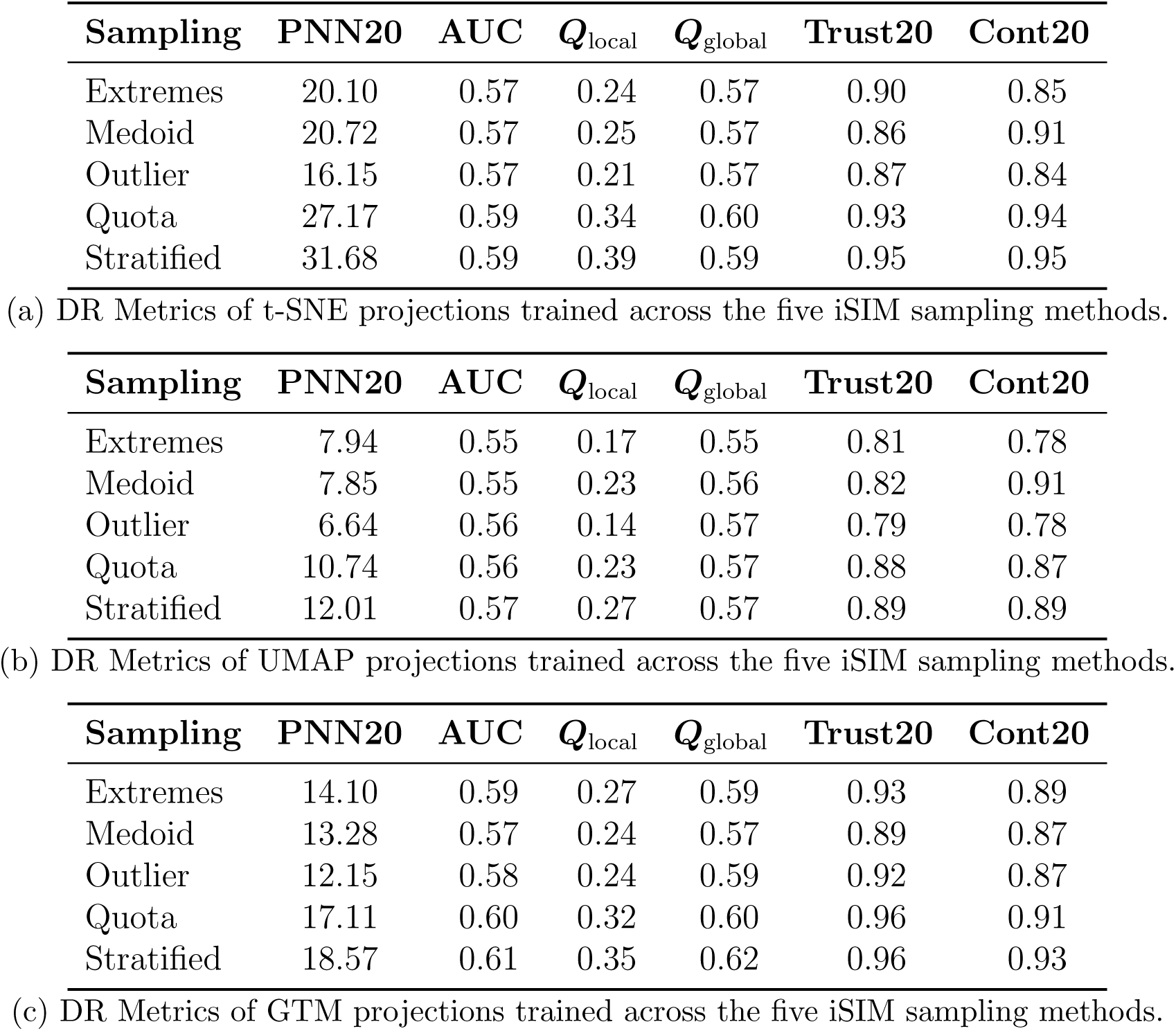
Embedding quality of neighborhood preservation metrics across the five sampling methods for (a) t-SNE, (b) UMAP and (c) GTM on the 50k molecule dataset. As before, the global preservation metrics do not show a large variation, local preservation generally show improvement in all three DR methods for quota and stratified samples.

The consistent underperformance of extremes, medoid and outlier sampling methods across all scales, methods and metrics points to a practically actionable conclusion: oversampled molecules from a particular region of the chemical space do not constitute a representative training set for manifold-learning-based DR methods regardless of dataset size. Stratified and quota sampling, by covering the full range of complementary similarity proportionally, reliably produce the most accurate hyperparameter selection at every scale tested, and should be the default choice for practitioners implementing this framework on new datasets. Finally, the stability of these trends across about two orders of magnitude from about 600-3600 molecules to 50000 molecules strongly suggests that the framework as a whole is likely to remain effective at larger scales, potentially extending to CHEMBL at its full size of several million compounds. A complete benchmarking of million molecule CHEMBL datasets is not practically feasible, since the calculation of these metrics stems from pairwise distance information which scales quadratically with dataset size, in both time and memory. The O(*N*) scaling of iSIM sampling ensures that diverse subset generation does not become a computational bottleneck even at very large N, and the consistency of the relative performance of sampling methods and DR models across scales provides a principled basis for expecting the approach to generalize without modification to the full chemical space.

### Conclusions

This work primarily focused on the effect of deterministic, diversity-driven undersampling strategies on the quality of non-linear DR for chemical space visualization from a scale of several hundreds to 50000 molecules. Training t-SNE, UMAP, and GTM on just 10% subsets generated by five iSIM complementary similarity sampling strategies, we find that the choice of training sample has a consistent and reproducible effect on projection quality, with stratified and quota sampling outperforming outlier-focused approaches across all methods and both scales on local preservation metrics including *P_NN_*, Trustworthiness and Continuity. Notably, the relative performance of DR methods is scale-dependent: GTM leads on local preservation for smaller datasets, while t-SNE emerges as the top performer at 50,000 molecules, a shift attributed to the larger absolute training pool available at scale, which enables more faithful perplexity optimization. UMAP remains competitive on global metrics but consistently underperforms on local neighborhood preservation, and PCA’s near-zero local preservation scores confirm that linear projections are categorically inadequate for local structure visualization in large, diverse chemical libraries.

The fundamental message of this work is that training non-linear DR models on small but carefully selected diverse subsets not only substantially reduces computational cost but can actively improve projection quality by guiding hyperparameter selection toward configurations that better capture the intrinsic local structure of chemical space. The O(N) scaling of iSIM sampling ensures that this approach remains computationally tractable as dataset size grows, and the consistency of results across two orders of magnitude of dataset size provides a principled basis for expecting the framework to generalize to the full CHEMBL database and beyond. Future work could explore wider hyperparameter grids for t-SNE and UMAP, the application of additional molecular representations beyond ECFP4 fingerprints, and the extension of this pipeline to even larger datasets approaching the scale of the complete drug-like chemical space.

## Acknowledgment

A.S., K.Z. and R.A.M.Q. thank the National Institute of General Medical Sciences of the National Institutes of Health for support under award number R35GM150620. A.S thanks Alexey Orlov for valuable insights on the neighborhood preservation metrics code.

## Notes

### Competing Interest Statement

The authors have declared no competing interest.

### Summary of Updates

Revised manuscript after referee reviews, more analyzes were included and discussion was improved

